# Bi-Alignments as Models of Incongruent Evolution of RNA Sequence and Structure

**DOI:** 10.1101/631606

**Authors:** Maria Waldl, Sebastian Will, Michael T. Wolfinger, Ivo L. Hofacker, Peter F. Stadler

**Affiliations:** University of Vienna, Faculty of Chemistry, Dept. of Theoretical Chemistry, Währingerstraße 17, 1090 Vienna, Austria. {,,, }; University of Vienna, Faculty of Computer Science, Research Group Bioinformatics and Computational Biology, Währingerstraße 29, 1090 Vienna, Austria; Dept. of Computer Science and Interdisciplinary Center for Bioinformatics, Leipzig University, Härtelstraße 16-18, 04109 Leipzig, Germany

**Keywords:** RNA secondary structure, RNA alignment, incongruent evolution

## Abstract

RNA molecules may experience independent selection pressures on their sequence and (secondary) structure. Structural features then may be preserved without maintaining their exact position along the sequence. In such cases, corresponding base pairs are no longer formed by homologous bases, leading to the incongruent evolutionary conservation of sequence and structure. In order to model this phenomenon, we introduce bi-alignments as a superposition of two alignments: one modeling sequence homology; the other, structural homology. We show that under natural assumptions on the scoring functions, bi-alignments form a special case of 4-way alignments, in which the incongruencies are measured as indels in the pairwise alignment of the two alignment copies. A preliminary survey of the `Rfam` database suggests that incongruent evolution of RNAs is not a very rare phenomenon.

**Availability:** Our software is freely available at *https://github.com/s-will/BiAlign*

## 1 Scientific Background

The secondary structure of many functional RNAs is well conserved over long evolutionary timescales. Paradigmatic examples include rRNAs, tRNAs, spliceosomal RNAs, small nucleolar RNAs, the precursors of miRNAs, many families of regulatory RNAs in bacteria, as well as some regulatory features in mRNAs, such as iron-responsive (IRE) or selenocystein insertion (SECIS) elements. The `Rfam` database [1] collects these RNAs and presents them as an alignment of sequences from different species annotated by a consensus secondary structure. In such families, the variation of the secondary structure is limited to small deviations from the consensus (additional or ommited base pairs). Even more stringently, the notion of a consensus structure implies that conserved base pairs are formed by pairs of homologous nucleotides.

If selection acts to preserve base pairs, then base pairs provide additional information of the homology of nucleotides. As a consequence, Sankoff’s algorithm [2] to simultaneously compute an alignment (of the nucleotide positions) and a consensus structure (by rewarding base pairs formed by pairs of aligned positions) results in an improvement of both the sequence alignment and the predicted secondary structure. Although this assumption of *congruent evolution* of sequence and structure is appealing (and has been very fruitful for modeling RNA families in the `Rfam` database), a partial survey of `Rfam` reveals several families that do not follow congruent evolution.

Fig.1 shows two alignments of the sequence and structure of two paralogous subfamilies of mir-30 precursors. The two families presumably are a product of the vertebrate-specific (2R) genome duplication and have evolved independently for the last 600 Myr. While the two alignments agree in the outer part of the stem loop structure, we observe that the structure alignment (bottom) slightly misaligns well matching sub-sequences, which are well-aligned by the sequence alignment (top), in order to properly align corresponding structure. As key observation, sequence and structure cannot be reconciled in this case. Insisting on matching common sequence patterns necessarily disrupts base pairs, while matching up the base pairs implies that the corresponding sequences appear “shifted” relative to each other.

In this contribution, we assume a very simple mechanism to bring about incongruencies between sequence and structure: as usual, we assume strong negative selection on both sequence and structure. However, we assume (1) that the selective pressures on sequence and structure are mechanistically independent, and (2) the exact position of the individual base pairs are less important than the overall ‘shape’ (e.g. the cloverleaf of a tRNA) of the secondary structure. In such a model, a stem may “move” by losing a base pair on one end and introducing a new base pair at the other end. While this example is still consistent with a consensus structure, in which all inner base pairs are conserved, it remarkably allows for more unusual evolutionary transitions.

In the simple example of evolutionary stem sliding, Fig. 2, the sequences of the two sides of a stem (or entire stem-loop) structure allow two different pairings with disjoint sets of base pairings but comparable energy. Single substitutions or indels may stabilize either one or the other structural alternative, leading to very similar sequences that also have very similar structures, while base pairs are no longer conserved for homologous nucleotides. As a consequence, the sequence alignment (describing *homologous* nucleotides) and the alignment of secondary structures (describing *analogous* base pairs) are incongruent. Stem sliding may explain the evolution of the mir-30 paralogs in Fig. 1: selective pressures at sequence level are dominated by stabilizing selection on the mature miRNA product, while pressures on the structure require only a sufficiently stable stem-loop structure to maintain Dicer processing, independent of the exact position of the mature product in the precursor hairpin.

**Figure 1:**
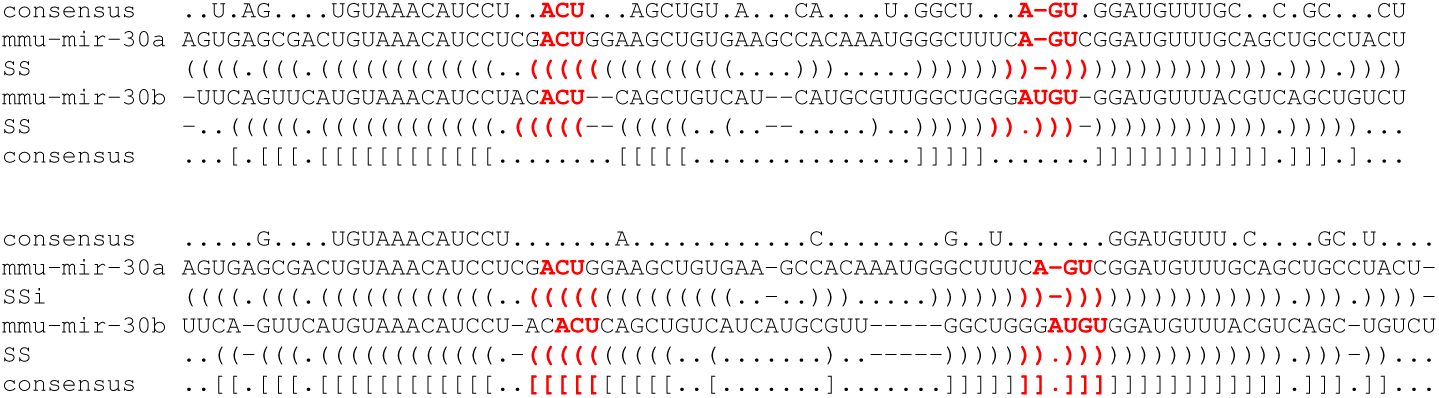
Alignments of the structure of mir-30a and mir-30b, two paralogous miRNAs that diverged since the genome duplications in the ancestor of the jawed vertebrates show evidence of incongruent evolution of sequence and structure. The sequence-based alignment (top) suggests that parts of the stem structure (missing in the consensus) have shifted relative to the sequence. The structure-based alignment (below) shows that the structures are nearly identical, while the underlying sequences is partially misaligned. The difficulties to reconcile the sequence and structure alignment (by only slight shifts) indicate the incongruence of the evolutionary history.

**Figure 2:**
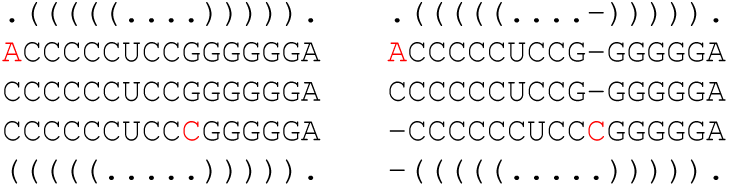
Evolutionary stem sliding. The two hairpins shown in “dot-parenthesis” notation have no base pair in common. The middle structure folds into both structures with similar energy, the mutants fix different alternatives.

The incongruence between sequence and structure alignment violates the assumptions underlying the consensus structure model: in a sequence based-alignment, no base pairs are conserved, and tools such as `RNAalifold` [3] that determine consensus structures are likely to fail. Conversely, structure-based alignments such as `RNAforester` [4] will produce nearly perfect structural alignments. These, however, match up non-homologous bases and thus grossly overpredict the turnover of base pairs. Combined sequence/structure alignments based on the Sankoff algorithm [2], such as `LocARNA` [5] or alternatives such as `cmfinder` [6] do not provide remedy, since these methods also assume congruent evolution of sequence and structure. This calls for a novel formal framework that makes it possible to capture incongruent evolutionary changes mathematically, which serves as a basis for developing algorithmic approaches to systematically study this phenomenon.

## 2 Theory

### Bi-Alignments

To resolve this conundrum, we embrace the idea that the evolution of sequence and structure is properly modeled by a pair of alignments that are related by evolutionary shifts between sequence and structure. Since the shift events couple the two alignments, they must be optimized multaneously, leading to a 4-way alignment for pairwise comparison. Naturally, this extends to 2*k*-way alignments for comparing *k* ≥ 2 sequence/structure pairs. Here we elaborate only the pairwise case for the sake of brevity. The dynamic programming algorithm outlined below solves the simultaneous alignment problem exactly for arbitrary sequence/structure pairs. Given two sequences *x* and *y* we consider two independent alignments 𝔸 and 𝔹. A plausible scoring model for their combination is to add their scores *s*_1_(𝔸) and *s*_2_(𝔹) and to introduce an additional term that penalizes shifts between them. To keep things simple at first, we model both scores based on indel scores and similarities between sequence positions. As elaborated later, this allows approximating the structure score. A shift occurs whenever 𝔸 and 𝔹 cannot agree on some operation. This naturally leads to a variant of 4-way alignments: we simply align *x* and *y* twice, with the first pair of rows corresponding to first scoring model and the second pair of rows corresponding to second scoring model.

The simplicity of both scores allows us to describe our efficient dynamic programming (DP) bi-alignment algorithm in the form of a regular grammar *A* → *A***c**, where c is one of the 15 different possible types of the last column in a 4-way alignment, ranging from 4 matches to deletions in three of the four rows [7]. Based on the column type **c**, one introduces shifts: if the first and second pair of entries agree, both alignments stay in sync (without any shift); otherwise, the alignments shift out of phase. Each shift by one position incurs an extra *shift penalty* Δ. The 15 cases and their cost are now be conveniently presented as columns of a table:

|  |  |  |  |  |  |  |  |  |  |  |  |  |  |  |  |
| --- | --- | --- | --- | --- | --- | --- | --- | --- | --- | --- | --- | --- | --- | --- | --- |
| $x$ | • | • | - | • | • | • | - | • | - | • | - | • | - | - | - |
| $y$ | • | - | • | • | • | - | • | • | • | - | - | - | • | - | - |
| $x$ | • | • | - | • | - | • | • | - | • | - | • | - | - | • | - |
| $y$ | • | - | • | - | • | • | • | - | - | • | • | - | - | - | • |
| $s(\mathbb{A})$ | $\mu$ | $\gamma$ | $\gamma$ | $\mu$ | $\mu$ | $\gamma$ | $\gamma$ | $\mu$ | $\gamma$ | $\gamma$ | 0 | $\gamma$ | $\gamma$ | 0 | 0 |
| $s(\mathbb{B})$ | $\mu$ | $\gamma$ | $\gamma$ | $\gamma$ | $\gamma$ | $\mu$ | $\mu$ | 0 | $\gamma$ | $\gamma$ | $\mu$ | 0 | 0 | $\gamma$ | $\gamma$ |
| shift | 0 | 0 | 0 | $\Delta$ | $\Delta$ | $\Delta$ | $\Delta$ | $2\Delta$ | $2\Delta$ | $2\Delta$ | $2\Delta$ | $\Delta$ | $\Delta$ | $\Delta$ | $\Delta$ |

The symbol *µ* indicates (mis)match scores for 𝔸 and 𝔹, which in practice differ for both scores and can be position-specific; *γ* and Δ respectively represent gap and shift scores. In three cases both alignments move “in sync” and thus incur no shift penalty. In four cases, however, the alignment moves out of sync simultaneously in both sequences, thus incurring twice the penalty. Consequently, bi-alignment reduces to a 4-way alignment with a sum-of-pair-like scoring scheme: For the two alignments of *x* and *y* (in the first and second pair of rows), the user-defined scoring models apply. Indels in the alignments of the sequence with themselves (rows 1&3 and 2&4) correspond to incongruencies and are penalized, while mismatches are ignored. The remaining two pairs of rows 1&4 and 2&3 are not scored at all.

### Limited shifting and complexity

Pairwise bi-alignment thus is a 4-way alignment problem, which can be solved with quartic time and memory by DP [8]. However, large numbers of shifts, i.e., large numbers of indels in the “self-alignments” of *x* and *y* are unlikely to be of interest in practice. Shifts can rather be strongly restricted, e.g. to 3 positions. This restriction can be easily realized in the algorithm by limiting index differences, such that one evaluates only a ‘band’ of the 4-dimension DP matrix. Consequently, one evaluates the recursions in quadratic time (w.r.t. the input length). It is tempting to simplify the 4-way alignment further by using either *x* or *y* as reference. In general, however, such a restriction can miss the optimal bi-alignment if it requires indels between the copies of *x* and the copies of *y*.

### Realistic Scoring of Structur

The (secondary) structure similarity of RNAs typically depends on similarities between corresponding (aligned) base pairs, which introduces a dependency between pairs of alignment columns. Moreover, in many cases the secondary structure of the RNAs is unknown, such that it must be inferred during the alignment. Both issues are addressed by computationally more complex algorithms often following the idea of Sankoff. As algorithmic short cut, one breaks the column dependencies and approximates structure similarity (resembling [9]) by a match similarity

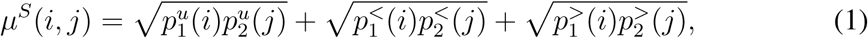

where the 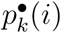 denote the respective probabilities that position *i* is unpaired (u), paired upstream (<), or downstream (>) in the ensemble of RNA k (*k* ∈ {1, 2}).

### Simultaneous Bi-alignment and Folding

The previous discussion naturally generalizes to a bi-alignment variant of Sankoff’s algorithm. It can be described by context-free grammar rules of the form *A* → *A***c**|*A*(*A*), where c denotes the extension of the 4-way alignment by a column as described for the simple bi-alignment model described above. The alternative production refers to a base pair in the consensus structure. More precisely, this production is of the form 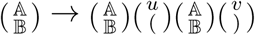 where the first coordinate refers to the sequence-based alignment and the second coordinate denotes the structural alignment. Here we only allow the insertion of a consensus base pair if both *x* and *y* support a base pair at the matching position. In the sequence part we may have 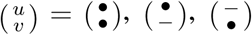, or 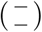, accounting for a total of 4 × 4 = 16 different cases. Both positions *u* and *v* independently respectively contribute *µ, γ* + Δ, *γ* + Δ, or 2Δ to the total score. In its most general form, this 4-way version of Sankoff’s algorithm has a space complexity of *O*(*n*^8^) and a time complexity of *O*(*n*^12^). Restricting the total shift between the two alignments to a small value, however, reduces the complexity to *O*(*n*^4^) and time complexity of *O*(*n*^6^). Notably, this can be improved further to *O*(*n*^4^) time and *O*(*n*^2^) space by transferring techniques from the Sankoff variant `LocARNA` [5].

### Generalization to Multiple Bi-Alignments and Poly-Alignments

The notion of bi-alignments can be generalized to a pair (𝔸, 𝔹) of multiple alignments of the sequences *x*^1^, *x*^2^, …, *x*^*k*^. The corresponding bi-alignment can be represented as 2*k*-way alignment, with a scoring function of the form 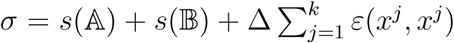,where *ε*(*x*^*j*^, *x*^*j*^) counts the indels between two copies of *y*^*j*^ in the 2*k*-way alignment. A natural choice of the scoring function is the sum-of-pair score scaled by 1*/k* to ensure that the contributions of the alignments and the shift penalties scale in the same way with *k*.

A different generalization are poly-alignments, which are superpositions of the three or more (pairwise) alignments. Again the problem can be represented as a multiple alignment problem with a special scoring function: In addition to the (pairwise) alignments 𝔸_*i*_, 1 ≤ *i* ≤ *𝓁*, the indels between the *𝓁* copies of the same sequence need to be accounted for. A natural score is the sum of indels between each pair of copies, normalized by 1*/*(*𝓁* − 1) to keep the balance between the *𝓁* pairwise alignment scores *s*(𝔸_*i*_) and 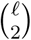 comparisons within the *𝓁* copies of each sequence.

Of course, these generalizations can be combined to multiple poly-alignments consisting of a superposition of *𝓁* multiple alignments of the same *N* sequences. The result is a *N* × *𝓁*-way alignment with a scoring scheme taking into account each multiple alignment as well as the indels between the copies of the same sequence.

## 3 Methods

### Candidate RNA families with Evolutionary Shifts of Sequence vs. Structure

The `Rfam` database provides a large set of curated alignments for structured RNAs. For many of these families the alignments have been created with an emphasis on the consensus structure. In order to identify `Rfam` families in which incongruent evolution may have played a rule, we compare `Rfam 14.1` seed alignments to their `MAFFT` [10] re-alignments. Employing a simple scoring function that considers the sequence positions observed in each column of the alignments, we determine cases, where the `Rfam` and `MAFFT` alignments strongly disagree. This can indicate deviance of sequence and structure evolution.

### Bi-alignment Enables Systematic Screening for Evolutionary Shifts

An increase of the combined sequence and structure similarity in the bi-alignment compared to the shift-free baseline serves as a good indication of evolutionary shift events. We systematically screened for such events by performing all pairwise alignments the set of RNA families selected as described above. Two types of alignments were computed using our bi-alignment algorithm: once allowing a shift by at most three positions (an intentionally conservative choice), and once by forbidding shifts (enforcing congruent evolution). In our preliminary study, we choose ad-hoc, but plausible paramters for assessing similarity: the sequence similarity in our bi-alignments is simply composed from scoring identical matching nucleotides with 100 (and mismatchs with 0). For assessing structure similarity, we distinguish two cases: for a priori unknown structure, we apply *µ*^*S*^ from Eq. 1; for known structures, we simply count the matched symbols in the dot-bracket structure strings. For weighting structure against sequence similarity, we multiply the structure similarity by 100. Finally, we score all indels with −200 and each shift with −250. Our script ranks the RNA pairs based on the observed score differences.

## 4 Results

### Implementation

We implemented our bi-alignment algorithm in Python 3 as an open source software tool. The implementation provides a convenient command line interface and alternatively can be integrated as a Python module; we utilize the latter in our screening pipeline. Both interfaces support full parametrization of the alignment scores and the maximum shift between the two alignments. Note that setting the maximum shift to zero provides a shift-free base line. To facilitate by-eye inspection, the tool can highlight conserved sequence and structure.

### Survey of Rfam for Incongruent Evolution of Sequence and Structure

To quickly find plausible candidate families which show incongruent evolution, we focused on `Rfam` families with small and medium-width seed alignments (≤ 10 sequences, ≤ 120 columns). This leaves us with 1181 of 3016 families in `Rfam 14.1`. Out of these we identify 709 cases where the Rfam alignment differs from the MAFFT realignments.

In a second step, we scrutinized all 10137 pairs of RNA sequences from the 709 alignments for indications of shift events. Each pair of sequences was aligned twice, with and without shifts. In 143 cases, the optimal bi-alignment exhibits at least one evolutionary shift event; these cases stem from 72 different Rfam families. Fig. 3 shows one example from this study, where an evolutionary shift event looks very plausible. Naturally, the number and significance of suggested shifts strongly depend on the similarity score (in particular, shift costs).

**Figure 3:**
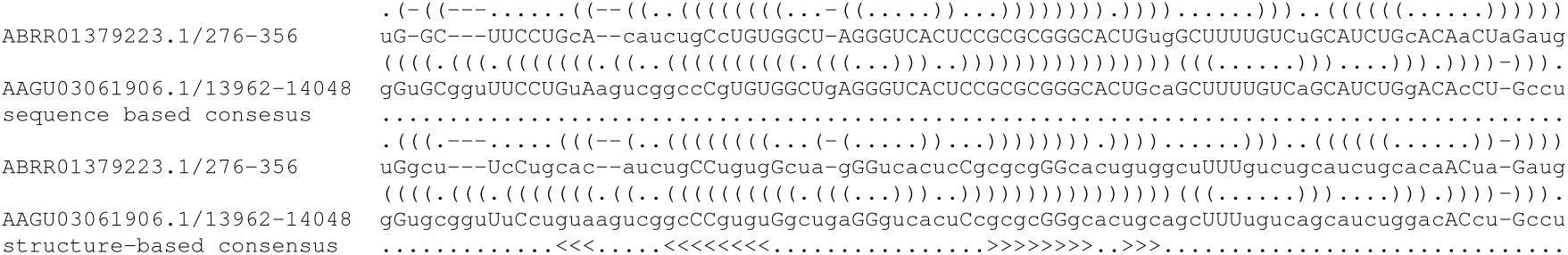
Shift alignment from conserved region 1 of the long non-coding RNA Six3os1 (Rfam family `RF02246`). For this instance, the alignment without shifts (not shown) does not align any base pairs correctly, since apparently the structure is shifted against the sequence by 1 position; shifts dramatically improve the alignment score. We show the annotated output of our program, where—for each of the two sub-alignments—the matched nucleotides are capitalized and matched base pairs appear in the consensus structure strings as balanced ‘<>’ pairs. The alignment is based on the mfe structures of the sequences, which are annotated above their corresponding sequences.

## 5 Conclusion

Incongruent evolution of sequence and structure cannot be captured by the existing RNA alignment methods, which focus on consensus structures. Instead of performing a single common alignment, sequence and structure alignment need to be represented separately to account for incongruencies. Bi-alignments appear to be a well-suited mathematical construction for this purpose. Here, we have shown that bi-alignments can be treated as 4-way (and in general 2*k*-way) alignments with a scoring function that evaluates the constituent alignments and the shifts between the two copies of the same input sequences. Limiting the total amount of shifts between sequence and structure alignment, the computational efforts exceeds the individual alignment problems only by a constant factor. Consequently, bi-alignments are not only of conceptual interest but are also computationally feasible.

## Acknowledgments

PFS is also affiliated with the Max Planck Institute for Mathematics in the Sciences in Leipzig, the Universidad Nacional de Colombia in Bogotá, and the Santa Fe Institute.

